# An Intra-Hypothalamic Pathway Modulating Body Temperature and Feeding

**DOI:** 10.1101/2024.04.16.589686

**Authors:** Sara Nencini, Jan Siemens

## Abstract

The intricate interplay between energy metabolism and body temperature regulation underscores the necessity of finely tuned mechanisms to maintain thermo-energetic homeostasis. Hot environments are known to suppress food intake and to reduce energy expenditure. However, the interplay between thermoregulatory and caloric-regulatory hypothalamic areas remains largely unexplored. In this study, we unveil two unconventional pathways originating from a subpopulation of genetically defined excitatory, leptin receptor-expressing POA neurons (VMPO^LepR^) that connect to the paraventricular nucleus of the hypothalamus (PVH) and the dorsomedial hypothalamic nucleus (DMH). Both, VMPO^LepR^→PVH and VMPO^LepR^→DMH connections, inhibit brown adipose tissue (BAT) thermogenesis and reduce body temperature; surprisingly, the VMPO^LepR^→PVH connection additionally exhibits the unique ability to suppress food intake and also promotes tail vasodilation. Our findings suggest that the excitatory VMPO^LepR^→PVH loop integrates temperature and caloric information to complement the canonical inhibitory arcuate nucleus (ARC)→PVH pathway. We propose that this novel pathway contributes to energy and temperature homeostasis in hot environments, offering new insights into previously unrecognized neuronal circuits orchestrating thermo-metabolic balance in response to environmental challenges.

## Introduction

Energy metabolism and body temperature are intricately linked. In mammals, body temperature is usually maintained within a narrow range, requiring finely tuned heat loss and heat gain mechanisms that partially depend on energy utilization. For example, cold ambient temperatures promote food consumption to meet energetic demands required for thermogenesis to keep the body warm. Oppositely, warm ambient temperatures suppress food intake and energy expenditure to maintain thermo-energetic homeostasis^1–3^.

The hypothalamus harbors neuronal centers that serve both regulatory processes, body temperature control and caloric balance. It has therefore been proposed that thermoregulatory and calory-regulatory hypothalamic areas interact to coordinate energetic and thermal optimums^4–7^ and several cell populations modulating both, feeding and body temperature have been characterized^8–11^. However, how different hypothalamic cell populations interact to govern the intricate thermo-metabolic balance and to resolve conflicting requirements and tradeoffs ––for example when scarcity of food prevents sustained thermogenesis in cold environments–– is only partially understood.

Temperature information from peripheral (and central) thermoreceptors are integrated in the hypothalamic preoptic area (POA) to regulate body temperature via peripheral effector organs such as brown adipose tissue, muscle and skin^5,12–14^.

The arcuate nucleus (ARC), on the other hand, is considered a main hub regulating energy homeostasis, and contains interoceptive neurons responding to caloric state to regulate feeding behavior^15,16^. Several studies demonstrate that projections from the ARC to the paraventricular nucleus of the hypothalamus (PVH) are crucial for regulating caloric homeostasis and feeding^17–20^. The PVH has so far not been considered a major area for body temperature regulation, except for a modulatory role of brown adipose tissue (BAT) thermogenesis^21,22^. On the other hand, reciprocal thermo-metabolic connections have been suggested to exist between the POA and the ARC^23,24^. Moreover, recent data suggests that neuronal pathways from the POA to the PVH can influence feeding behavior^23^.

Circulating leptin serves as an indicator of energy status^25^ and the hormone also plays a major role in body temperature regulation^26^. Interestingly, leptin receptor-positive neurons in the rostral part of the POA, the so-called ventromedial POA (VMPO), are warm-responsive neurons activated by ambient temperature increases that modulate both, body temperature and food consumption^11,27^. Specifically, activating these neurons *in vivo* triggers profound hypothermia and inhibits food intake. However, the neural pathways that these thermoregulatory neurons utilize to precisely regulate food intake in accordance with thermo-metabolic balance and body temperature stability remain elusive.

We here describe a pathway that originates from a subpopulation of excitatory leptin receptor-expressing POA neurons (VMPO^LepR-Glut^ neurons) that inhibit feeding and diminish energy expenditure. Excitatory VMPO^LepR^ neurons do not exclusively target the PVH and we also find connections to the Periaqueductal Grey (PAG), the arcuate nucleus (ARC) and Dorsomedial Hypothalamus (DMH), the latter also mediating heat loss responses. However, monosynaptic connections of VMPO^LepR-Glut^ neurons to the PVH appear to be unique in that they orchestrate inhibition of food intake and energy expenditure while simultaneously also mediating heat loss mechanisms.

We propose that this excitatory VMPO^LepR^èPVH loop integrates temperature information to complement and fine-tune the well-established inhibitory ARCèPVH pathway, thereby conjointly achieving energy and temperature homeostasis in hot environments.

## Results

### Excitatory VMPO^LepR^ neurons promote hypothermia and inhibit food intake

To investigate the impact of VMPO^LepR^ neuron activity on body temperature, we photostimulated LepR cell bodies using optic fibers implanted above the VMPO. Light stimulation at frequencies of 5 Hz or higher resulted in a drop in body– and brown adipose tissue (BAT) temperature, a decrease in overall ambulatory activity, and a suppression of food intake in mice expressing channelrhodopsin-2 (ChR2) in VMPO^LepR^. (Fig. 1). These findings corroborate previous observations from chemogenetic manipulation experiments^11^. Additionally, VMPO^LepR^ neuron activation transiently triggered cutaneous vasodilation (Fig. 1C). These thermoregulatory and energy-metabolic responses were dependent on ChR2 expression and were not observed in (mCherry-expressing) control animals (Fig. 1B). Furthermore, these responses appeared to be independent of the hypothalamic–pituitary–adrenal axis, as corticosterone levels remained at baseline levels (Fig. 1G).

**Figure 1.**
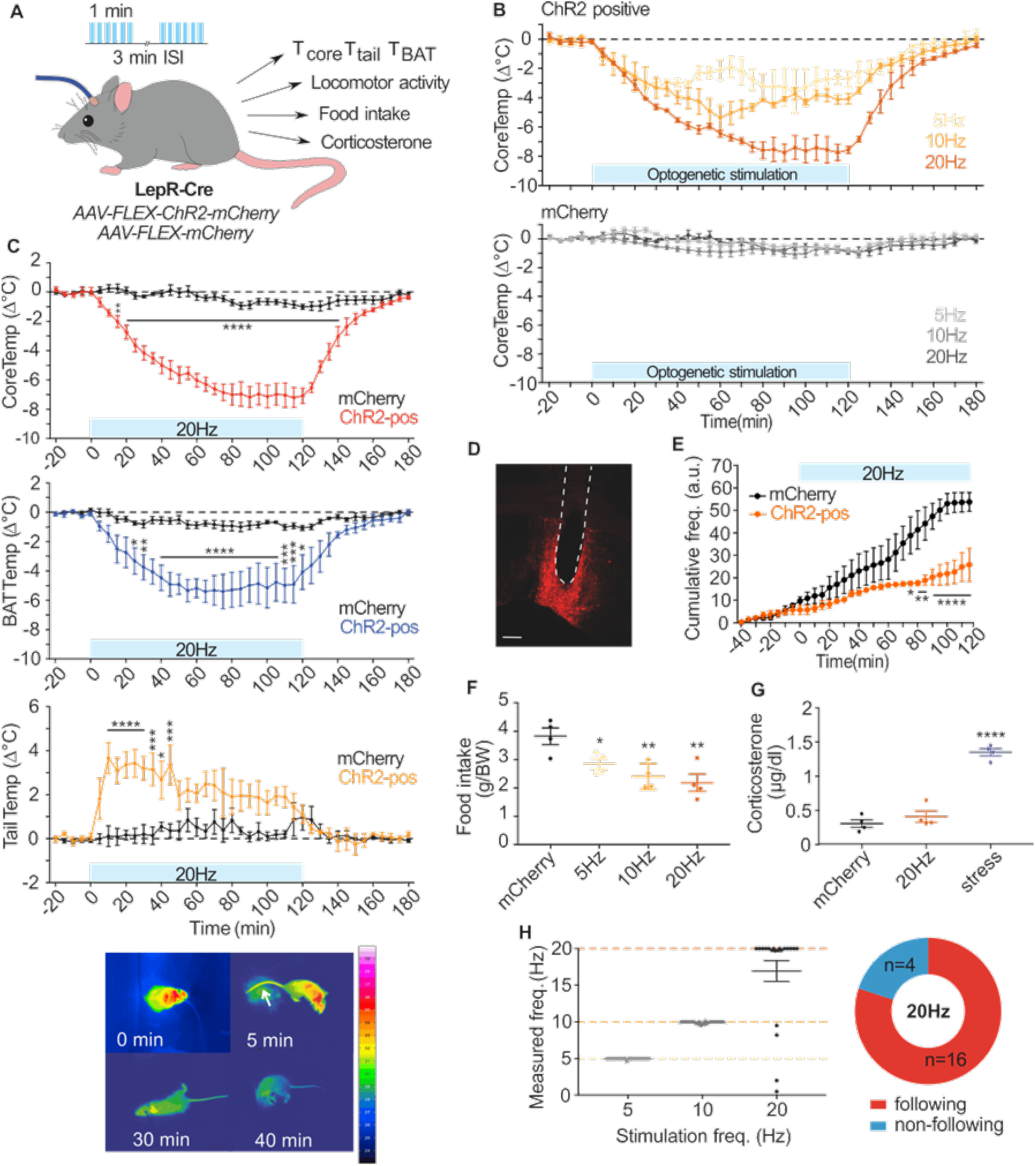
– Optogenetic stimulation of POA^LepR^ neurons mediates body cooling and a reduction in energy intake/expenditure by engaging multiple mechanisms. **A.** Schematic representation of the optogenetic experiment. Optogenetic stimulation was applied unilaterally using pulses of 10ms duration (473nm, max 6mW) with varying stimulation frequencies (5Hz, 10Hz, 20Hz) for 1 min, followed by a 3-minute inter-stimulation-interval (ISI). Animals (ChR2-positive and Chr2-negative mCherry controls) were analyzed for changes in core body temperature (T_core_), BAT (T_BAT_) and tail temperature (T_Tail_), locomotor activity, food intake and corticosterone levels. **B.** Pooled data showing that optogenetic stimulation of VMPO LepR cells reduced body core temperature in ChR2 injected animals (top panel) but not in mCherry controls (bottom panel; n = 4 mice for each group) at all stimulation frequencies. The maximum decrease in CoreTemp was observed towards the end (80-120 min) of a 20Hz stimulation protocol. Blue bar represents time the light pulse-protocol was applied. **C.** Comparison between ChR2-positive and ChR2-negative animals upon optogenetic stimulation. 20Hz pulse stimulation (blue bar) decreased body core temperature (upper panel) as well as BAT temperature (middle panel) while transiently increasing tail temperature (lower panel), specifically in ChR2-positive animals. Two-way ANOVA (effect of stimulation x time), p < 0.0001; Dunnett’s multiple comparison test: *p < 0.05, **p < 0.01, ***p < 0.001, ****p < 0.0001. Bottom panel: Representative thermographic images obtained while optogenetically activating LepR neurons (20Hz). Tail temperature increased 5min after beginning the light pulse protocol (arrow). **D.** Representative histological image showing optical fiber placement (white dotted line) and ChR2-mCherry unilateral injection in LepR cells in the VMPO area. Scale bar: 200µm. **E.** Mean cumulative locomotor activity [arbitrary units (A.U.)] of ChR2-positive and mCherry control mice. Optogenetic stimulation reduced the locomotor activity. Two-way ANOVA, (effect of stimulation x time), p < 0.0001, Dunnett’s multiple comparison test: *p < 0.05, **p < 0.01, ***p < 0.001, (N=3 mice per group). **F.** Food intake was significantly suppressed in ChR2-positive animals upon light activation, compared to mCherry controls (n = 4 mice for each group) with light stimulation performed overnight. One-way ANOVA; Dunnett’s multiple comparison test: *p < 0.05, **p < 0.01 vs mCherry group. **G.** Overnight optogenetic stimulation did not increase corticosterone content detected in feces of either mCherry or ChR2-positive animals. Animals that underwent a stress-restraining paradigm were used as positive controls for comparison. One-way ANOVA; Dunnett’s multiple comparison test: ****p < 0.0001 vs mCherry group. **H.** Left panel: responses of POA LepR neurons to 60 pulses of blue light at 5, 10 and 20Hz in electrophysiological *ex vivo* brain slice recordings. Right panel: Pie chart representing the percentage of neurons that were able (following) or not able (non-following) to induce action potentials at the same frequency as the indicated light stimulation frequency. Note that 4 out of 20 recorded LepR cells failed to consistently generate action potentials at the higher stimulation frequency. Data in B, C, E, F, G and H are shown as mean ± SEM.

Recent results suggested that excitatory POA neurons play a more prominent role in modulating body temperature than previously anticipated^12,27,28^. Warm-responsive VMPO^LepR^ neurons encompass both, excitatory (VGLUT2-positive) and inhibitory (VGAT-positive) neurons^11^. We therefore used an intersectional approach, employing LepR-Cre;vgat-FlpO mice in combination with adeno-associated viruses (AAV) that enables ChR2 expression in a Cre– and FlpO-dependent manner, to assess whether one of the two LepR-positive subgroups had a more pronounced effect on feeding, thermogenesis and tail vasodilation (Fig. 2A-C). With this approach we did not target the excitatory (VGLUT2-positive) population directly but activated either inhibitory (vgat-FlpO-on) or non-inhibitory (vgat-FlpO-off) VMPO^LepR^ neurons, presuming that the vgat-FlpO-off VMPO^LepR^ population is largely comprised of excitatory neurons. For simplicity we from here on refer to these populations as VMPO^LepR-inh^ (GABAergic) and VMPO^LepR-exc^ (glutamatergic) neurons.

**Figure 2.**
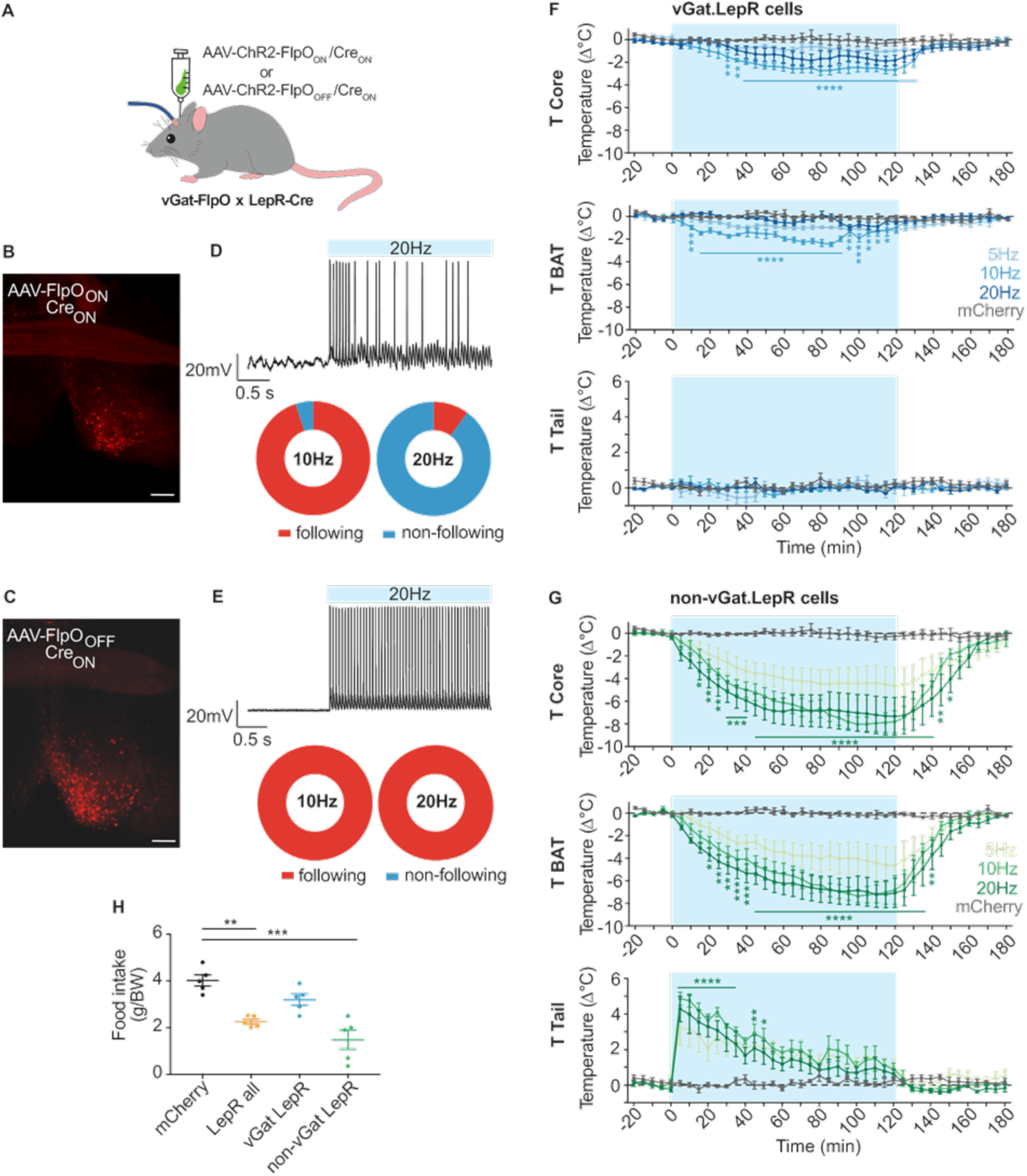
– Excitatory POA^LepR^ neurons drive hypothermia and reduction in food intake. **A**. Schematic indicating the strategy to selectively activate inhibitory (vGat-positive) or excitatory (vGAT-negative, non-inhibitory) VMPO^LepR^ neurons using an intersectional adeno-associated virus (AAV) genetic strategy to express ChR2 in FlpO– and/or Cre recombinase dependent fashion using vGAT-FlpO; LepR-Cre mice. **B** and **C.** Representative histological images showing the expression of rAAV(DJ)-Cre(ON)/FlpO(ON)-ChR2 and rAAV(DJ)-Cre(ON)/FlpO(OFF)-ChR2 virus, respectively in LepR-cre;vGat-FlpO mice. Scale bar 200µm. **D.** and **E.** Upper panels: Example trace of action potential response evoked by blue light stimulation at 20Hz stimulation frequency in FlpO_ON_, Cre_ON_ brain slices (D) or FlpO_OFF_, Cre_ON_ brain slices (E). Blue bar represents light pulse. Lower panels: Pie charts showing the percentage of neurons that were able (following) or not able (non-following) to induce action potentials at the same frequency as the indicated light stimulation frequency using 10 or 20Hz stimulation frequency in electrophysiological *ex vivo* brain slice recordings. Note that stimulation at 20Hz (in contrast to 10Hz stimulation frequency) failed to consistently elicit action potentials in the majority of vGat-positive LepR neuronal population. **F.** Pooled data showing that optogenetic stimulation of VMPO vGat-positive LepR cells slightly but significantly reduced body core (upper panel) and BAT temperature (middle panel) in ChR2 injected animals, but not in mCherry controls at 10Hz stimulation frequency (N = 5 mice for each group). An increase in tail temperature (lower panel) was not observed upon stimulation of this neuronal subgroup. Two-way ANOVA (effect of stimulation x time), p < 0.0001 for T core and T BAT; Dunnett’s multiple comparison test (indicated for 10Hz stimulation frequency only): *p < 0.05, **p < 0.01, ***p < 0.001, ****p < 0.0001. **G.** Pooled data showing that optogenetic stimulation of VMPO vGat-negative LepR cells reduced body core (upper panel) and BAT (middle panel) temperature in ChR2 injected animals, but not in mCherry controls (below) at all stimulation frequencies (n = 5 mice for each group). ChR2 activation of this neuronal subgroup also transiently elevated tail temperature (lower panel). Two-way ANOVA (effect of stimulation x time), p < 0.0001 for all graphs; Dunnett’s multiple comparison test (represented for the 10Hz stimulation frequency): *p < 0.05, **p < 0.01, ***p < 0.001, ****p < 0.0001. Data in F, G and H are shown as mean ± SEM. **H.** Food intake was significantly suppressed only when non-vGat LepR cells were light activated at 10Hz compared to mCherry controls and to a similar extent as for the whole LepR population (n = 4 mice for each group). One-way ANOVA; Dunnett’s multiple comparison test: **p < 0.01, ***p < 0.001.

We first confirmed in *ex vivo* brain slice preparations that optical stimulation, using different light pulse frequencies (5Hz, 10Hz and 20Hz), triggers corresponding action-potential frequencies in ChR2-expressing neurons. This experiment showed that both VMPO^LepR-inh^ and VMPO^LepR-exc^ neurons could sustain stimulation frequencies of up to 10Hz, but that inhibitory neurons did not faithfully recapitulate 20Hz optical stimulation frequencies (Fig. 1H and 2D, E). These results are in agreement with previous findings showing that galanin-positive POA neurons, which are predominantly inhibitory neurons, similarly failed to respond to higher frequencies^29^.

At optic stimulation frequencies that are compatible with triggering tonic action potentials in excitatory and inhibitory neurons, we find that VMPO^LepR-exc^ neurons inhibited BAT thermogenesis, transiently activated cutaneous vasodilation and suppressed food intake. Optic activation of VMPO^LepR-inh^ neurons only resulted in a small but non-significant drop in food intake and less pronounced responses on thermogenesis and tail vasodilation (Fig. 2F, G). These results agree with previous studies, targeting preoptic VGLUT2-expressing excitatory neurons non-selectively^11,23,30^.

### A VMPO^LepR^èPVH pathway suppresses feeding and promotes body cooling

We next asked which target areas downstream of VMPO^LepR^ neurons mediate thermoregulatory and feeding responses and whether the two functionalities are controlled by separate pathways. The POA is connected to several hypothalamic and extra hypothalamic brain areas with POA fibers prominently innervating the ventrolateral septal area (LSV), the bed nucleus of the stria vascularis (BNST), the ARC, the paraventricular thalamus (PVT), the medial habenula (mHB), the ventrolateral Periaqueductal Grey (vlPAG), Dorsomedial Hypothalamus (DMH) and the Paraventricular Hypothalamus (PVH)^31^. All of these regions have been implicated in the control of feeding^32–34^, while the DMH and to a lesser extent also vlPAG have been additionally implicated in thermoregulation^5,35^. We found fiber terminals of VMPO^LepR^ neurons to intensely innervate the PVH and DMH, and to a lesser extent we observed fiber labelling also in the PAG (Fig. 3A, B). Particularly, the PVH was robustly and strongly innervated by VMPO^LepR^ neuron fibers (Supplementary Fig. 1); only very few fibers innervated the ARC (data not shown). We therefore focused on the PVH, DMH, and PAG as potential relay stations for thermoregulatory and feeding-regulatory VMPO^LepR^ neuron projections.

**Figure 3.**
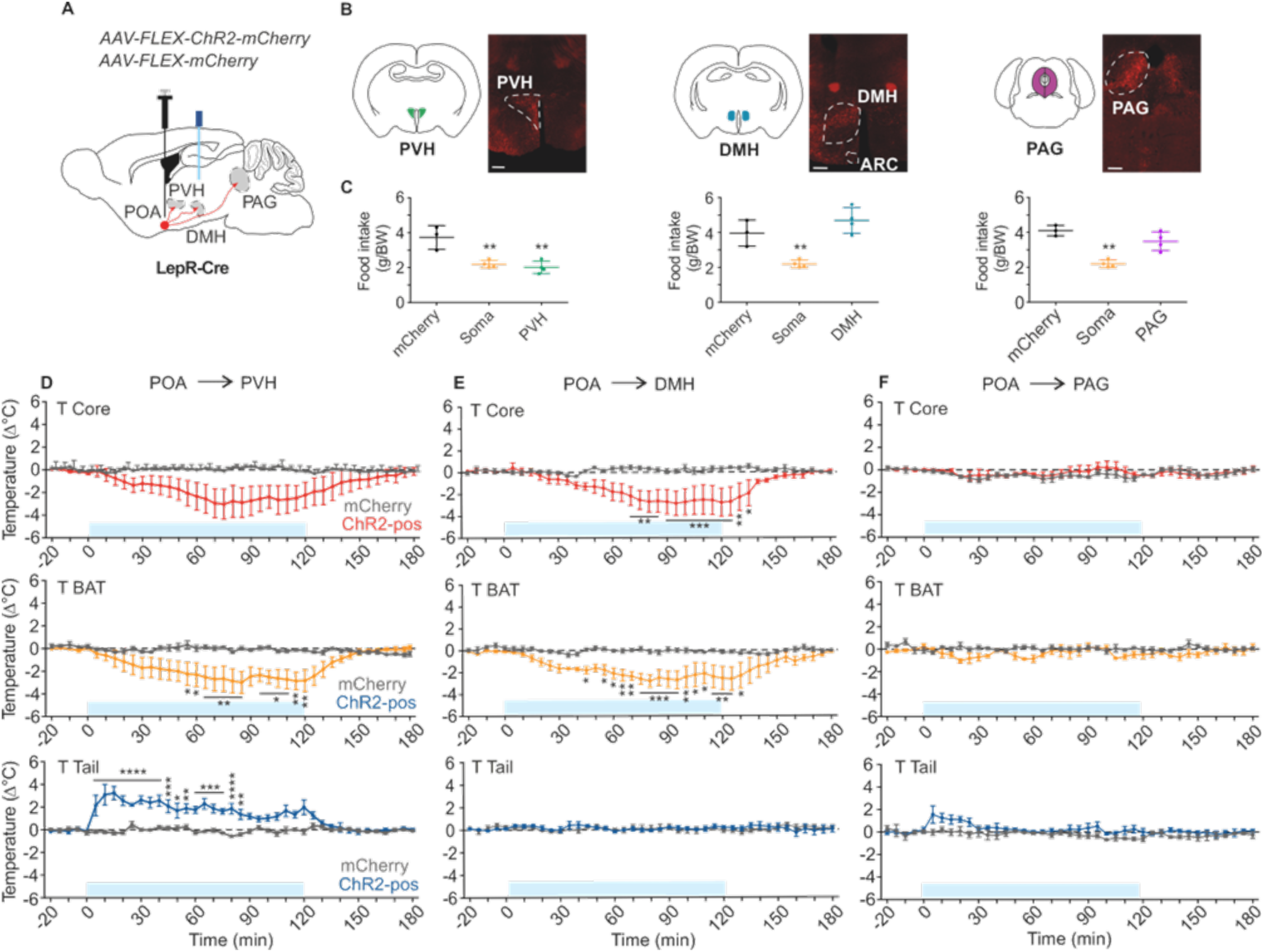
– Anterograde tracing and optogenetic stimulation of downstream LepR thermoregulatory circuit components. **A**. Schematic summarizing the strategy used for optogenetic activation of LepR terminals in three downstream projection areas. The major POA^LepR^ neuron projection sites are indicated: VMH – ventromedial hypothalamus, ARC – arcuate nucleus, PAG – periaqueductal grey. Control groups were injected with mCherry-AAV and stimulated at the level of LepR projection areas. **B.** Schematic representation of coronal sections and corresponding histological images showing ChR2-mCherry fluorescence in the LepR cell terminals innervating PVH, DMH and PAG (white dotted areas) as indicated. Scale bar: 200µm. **C.** Food consumption normalized to 30gr body weight during overnight optogenetic stimulation of VMPOLepR somas or PVH and DMH projection sites. Note that food intake was significantly suppressed when LepR to PVH (but not to DMH and PAG) projections were light activated at 10Hz. One-way ANOVA, **p < 0.01. (N = 4 mice for soma, PVH, DMH and PAG groups; N = 3 mice for control group; note that the “soma” data (somatic activation) is partially overlapping with data shown in Fig. 2H). **D.** Optogenetic stimulation at 10Hz of VMPO^LepR^ neuron terminals in the PVH of ChR2-positive animals (N = 5 mice) induced a decrease in body core (upper panel) and BAT (middle panel) temperature, as well as an increase in tail temperature (bottom panel), compared to mCherry controls (N = 3 mice). Two-way ANOVA (effect of stimulation x time), p=0.0074 for Tcore, and p < 0.0001 for BAT and Tail temperature; Dunnett’s multiple comparison test: *p < 0.05, **p < 0.01, ***p < 0.001, ****p < 0.0001. **E.** Optogenetic stimulation at 10Hz of VMPO^LepR^ neuron terminals in the DMH of ChR2-positive animals (N = 4 mice) induced a decrease in body core (upper panel) and BAT (middle panel) temperature, but not an increase in tail temperature (bottom panel), compared to mCherry controls (N = 3 mice). Two-way ANOVA (effect of stimulation x time), p < 0.0001 for T core and BAT; Dunnett’s multiple comparison test: *p < 0.05, **p < 0.01, ***p < 0.001, ****p < 0.0001. **F.** In contrast, optogenetic stimulation at 10Hz of VMPO^LepR^ neuron terminals in the PAG of ChR2-positive animals (N = 4 mice) did not induce any change in body core and peripheral effector organ temperature (T_BAT_ and T_Tail_), similar to mCherry controls (N = 3 mice). Data in C, D, E and F are shown as mean ± SEM.

To assess the role of VMPO^LepR^ projections innervating these 3 target areas, we again used an optogenetic approach by virally expressing ChR2 in VMPO^LepR^ neurons and directing the light source to the different target areas in order to selectively simulate fiber terminals in the DMH, PAG or PVH (Fig. 3A).

While activating projections innervating the PAG did not have any influence on body temperature, those innervating the DMH and PVH both decreased body temperature, likely via inhibiting BAT thermogenesis (Fig. 3D-F). Projections to PVH, but not those innervating the DMH, additionally mediated transient tail vasodilation (Fig. 3D-F). Strikingly, only activating the VMPO^LepR^→PVH pathway significantly reduced food intake, similar to activating the VMPO^LepR^ cell bodies (somatic activation) (Fig. 3C).

Because projections to the DMH and PVH had overlapping (but not identical) functionalities in the context of thermoregulation, while only PVH projections suppressed food intake, we assessed to what extent VMPO^LepR^ neurons innervating the PVH and DMH overlap. Injecting retroAAV particles into the DMH and PVH permitted the infection of fiber terminals to label upstream VMPO^LepR^ neurons with Cre-recombinase dependent red and green fluorophores, respectively. We found that only a small fraction of neurons overlapped, suggesting that collateralization to both areas is limited and the larger fraction of VMPO^LepR^ neurons appeared to project to either the PVH or the DMH (Supplementary Fig. 2).

### Excitatory but not Inhibitory VMPO^LepR^ projections to the PVH and DMH mediate thermoregulatory and feeding responses

Next, we tested whether excitatory (VGLUT2-positive) connections from the VMPO to the PVH could mediate the observed thermoregulatory effects and therefore injected AAV expressing ChR2 in a Cre-dependent manner in the VMPO of Vglut2-Cre mice. Optic stimulation of neuronal projections in the PVH triggered a T_BAT_ and T_core_ temperature drop and an increase in T_tail_ (Supplementary Fig. 3), similar to VMPO^LepR^→PVH projections (Fig. 3). These results are different to previous results that did not show any thermoregulatory responses when indiscriminatorily activating all VGLUT2-positive VMPO→PVH projections^23^.

We then utilized the intersectional approach (Fig. 2A) to further investigate the (excitatory or inhibitory) nature of the projections to PVH and DMH. Optogenetically activating excitatory ––but not inhibitory–– VMPO^LepR^ projections to PVH, and to a lesser extent DMH, recapitulated BAT inhibition and hypothermia induction (Fig. 4A-D). Activating the PVH projections also induced transient tail vasodilation (Fig. 4A). Inhibition of food intake was most robustly observed when activating excitatory VMPO^LepR^ projections to the PVH (Fig. 4E, F), a slight inhibition of food intake was also seen when activating terminals innervating the DMH.

**Figure 4.**
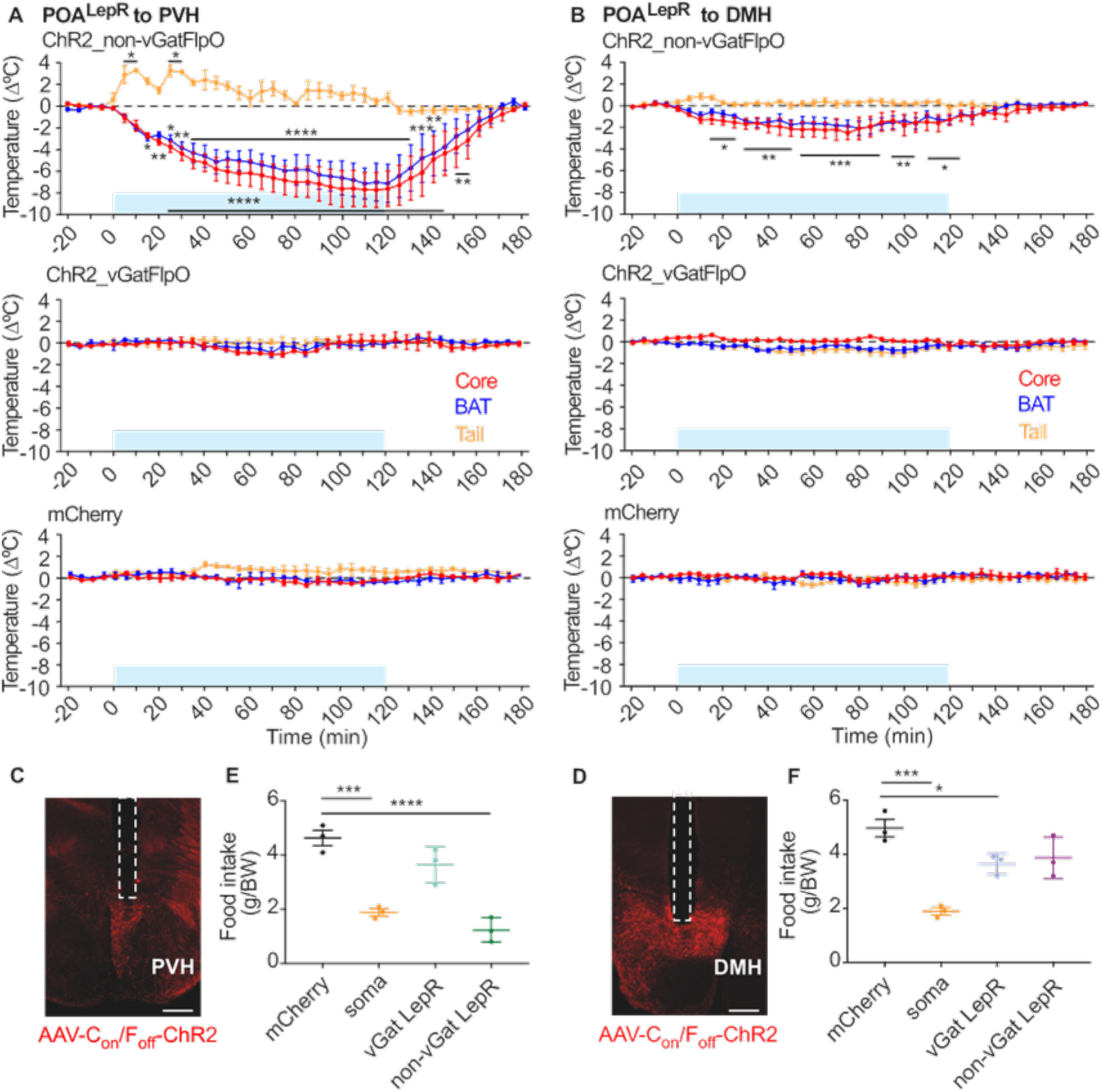
– vGAT-negative (excitatory) POA^LepR^->PVH neuron projections inhibit energy expenditure and food intake. **A**. Optogenetic stimulation of VMPO^LepR^ neuron terminals in the PVH at 10Hz of non-vGat LepR neurons (upper panel), but not of vGat LepR neurons (middle panel) induced a decrease in body core and BAT temperatures, as well as an increase in tail temperature, compared to mCherry controls (bottom panel) (N = 3 mice for each group). One-Way ANOVA comparing baseline (20 to 0 min) to stimulation/post-stimulation periods (0 to 180 min) separated for each temperature measurement paradigm (T_core_, red; T_BAT_, blue; T_tail_ yellow) **B.** Optogenetic stimulation of VMPO^LepR^ neuron terminals in the DMH at 10Hz of non-vGat LepR neurons (upper panel) resulted in a small but significant decrease in body core and BAT temperature. Conversely, optogenetic stimulation of vGat LepR neurons (middle panel), similarly to the mCherry control group (bottom), did not produce any temperature changes (N = 4 mice for each group). **C.** and **D.** Representative histological image showing the projection areas of vGat_OFF_ / LepR_ON_ neurons (indicated in red, ChR2-mCherry) and the optical fiber tract implantation site (indicated by dotted white line) in the PVH (C) and the DMH (D), respectively. Scale bar 200µm. **E.** and **F.** Food consumption normalized to 30gr body weight during overnight optogenetic stimulation of VMPO^LepR^ somas or PVH and DMH projection sites as indicated. Note that food intake was significantly suppressed when vGAT-negative (excitatory) VMPO^LepR^ neuron projections to PVH (but not to DMH) were light activated at 10Hz; One-way ANOVA (N = 3 mice for each group). Data in A, B, E and F are shown as mean ± SEM.*p < 0.05, **p < 0.01, ***p < 0.001, ****p < 0.0001.

Collectively, these results suggest that independent parallel pathways to the PVH and DMH, originating from excitatory VMPO^LepR^ neurons, regulate energy expenditure by inhibiting BAT thermogenesis and tail vasodilation, while food intake appears to be more selectively regulated via VMPO^LepR-exc^èPVH projections.

## Discussion

Several recent studies have implicated hypothalamic preoptic neurons in the suppression of energy expenditure and body temperature in mice and activation of some of those neuronal populations also inhibited energy (food) intake^11,27,30,31,36^. At the extreme, activity of these neurons results in a long-lasting “low energy state” of mice^37–39^, similar to the state of torpor^40^. Therefore, these neurons have been dubbed quiescence-inducing neurons or, based on the diverse “marker” genes they express, QPLOT neurons (which stands for <u>Q</u>rfp, <u>P</u>tger3, <u>L</u>epR, <u>O</u>pn5, and <u>T</u>acr3 mRNA transcripts that these VMPO neurons co-express)^41^.

However, the downstream neuronal pathways activated by these neurons are only incompletely understood.

We here described VMPO^LepR^èPVH and VMPO^LepR^èDMH connections that conjointly ––and likely via parallel and independent pathways–– inhibit BAT thermogenesis and reduce body temperature, with the VMPO^LepR^èPVH connection additionally promoting tail vasodilation and inhibiting food intake.

Recently it was shown that activation of excitatory (VGLUT2-positive) rostral POA neurons projecting to the PVH can inhibit food intake^23^. However, the authors of this study did not observe an effect on body/rectal temperature. This perceived difference compared to our results (showing that excitatory VMPO^LepR^ neurons projecting to the PVH engage in both, inhibiting food intake and inhibiting energy expenditure/mediating hypothermia) could be technical in nature, since in the other study rectal temperature was measured (and not intra-abdominal T_core_, using a telemetric temperature probe, as in our study); stimulating the presumably more heterogenous neuronal VGLUT2-positive fiber-terminal population in the PVH, compared to VMPO^LepR-exc^ fibers terminating in the PVH appears to have a less robust effect on T_core_ and T_BAT_ (compare Supplementary Fig. 3 and Fig. 4A) that could be missed when recording (less-sensitive) rectal temperatures. Additionally, the anatomical position of the VGLUT2 population targeted by Qian et al. might not be exactly the same compared to the relatively discretely positioned population of VMPO^LepR^ neurons^11^ that is examined here. Extensive local and distal connections emanate from the functionally heterogenous rostral POA/VMPO that could have opposing effects, thereby providing a potential explanation for the partially divergent results. In this regard it is interesting to note that we observed varying magnitudes of body temperature responses, depending on whether we optogenetically stimulated fiber terminals versus neuron-cell bodies (somas). One potential explanation for this difference is that VMPO^LepR^ projections to PVH and DMH may interact synergistically, so that the conjoint activation of both subgroups of VMPO^LepR^ together has the strongest effect. However ––and considering that fiber terminals innervating the PVH/DMH in experiments shown in Fig. 3 are both excitatory and inhibitory–– it is conceivable that the VMPO^LepR^ inhibitory connections might partially counteract the thermo-metabolic effects. Indeed, when only excitatory VMPO^LepR^ terminals are engaged by light stimulation of the PVH (Fig. 4A), the effect more closely resembles the response observed when VMPO^LepR^ cell somas are stimulated, arguing for a potential opposing effect by inhibitory terminals.

Interestingly, a sympatho-inhibitory population has been postulated to reside in the PVH that, when activated, inhibits BAT thermogenesis^21^. This elusive group of PVH neurons has been further characterized to receive inhibitory input from RIP-Cre neurons in the arcuate nucleus^22^. RIP neuron activity thereby increases BAT thermogenesis by inhibiting tonic activity of the PVH population. The origin of the tonic activity of these neurons is unknown as is their molecular identity. We hypothesize that VMPO^LepR^ neurons contribute to the activity of these sympatho-inhibitory PVH neurons. We further speculate that VMPO^LepR^ neurons also innervate MCR4-positive and/or PDYN-positive PVH neurons known to inhibit food intake^20^. Any PVH population that could mediate transient tail vasodilation, is to the best of our knowledge, currently unknown.

Our study indicates that, alongside the established ARC to PVH pathway, there exists an additional pathway from the POA to PVH that decreases food intake and reduces body temperature. The PVH contains different subpopulations of neurons that selectively modulate either food intake or energy expenditure. Pathways modulating either food intake or energy expenditure separately have been extensively described for the PVH^8,22,32,42,43^. However, a PVH pathway integrating both functions has, to the best of our knowledge, not been described.

Considering potential physiological rationales of this bifunctional pathway raises intriguing questions: at first glance, inhibiting energy expenditure and food intake simultaneously may seem counterintuitive: when energy is low, there should be a strong drive to take up more energetic fuel/food. When would such a low energy state be physiologically relevant and when would this VMPO^LepR^ neuronal network be activated under physiological conditions?

As mentioned above, the state of torpor appears to be an extreme thermo-metabolic condition that engages these neurons ^37–39,41^.

Given that the VMPO neurons we studied here are primarily marked by the leptin receptor, it is reasonable to assume that leptin may modulate the observed effects. Intriguingly, leptin has been shown to regulate torpor and if injected into fasted animals can reverse torpor^44^. Since the increase in neuronal activity of these QPLOT neurons can drive animals into a torpor-like state, it seems reasonable to assume that leptin should inhibit their neuronal activity to reverse the effect. Indeed, a subpopulation of VMPO^LepR^ neurons appears to be inhibited by leptin (while other VMPO^LepR^ neurons can be excited by leptin application, at least under baseline conditions)^45^. Interestingly Yu et al. find that *in vivo* leptin application into the POA/VMPO appears to influence energy expenditure mostly (or exclusively) when the animals are fasted^45^ ––the same condition/state required to induce torpor. Thus, it is possible that leptin’s capacity to exert regulatory control over these neurons is state dependent and may be at a maximum in a fasted state.

Whether fasting or other internal states alter the (excitatory vs. inhibitory) modulatory capacity and/or sensitivity of leptin on VMPO^LepR^ neurons is, to best of our knowledge, currently unknown.

There is an additional physiological scenario which dictates that both, energy expenditure and energy intake (food intake), are reduced to a minimum: in warm/hot environments when animals need to prevent overheating.

Intriguingly, VMPO^LepR^ neurons have been shown to respond to peripheral warmth/heat stimuli. Exposure to a warm/hot environment for an extended period of time is known to inhibit food intake, suppress energy expenditure and will result in recruiting heat loss responses^1^, all these aspects together will prevent overheating and are observed when the VMPO^LepR^→PVH circuit is activated.

It is therefore plausible that another primary function of VMPO^LepR^ (QPLOT) neurons (next to torpor induction) is to permit animals to adapt and survive in warm environments.

Future work will have to uncover the physiological relevance and downstream pathways of VMPO^LepR^/QPLOT neurons not only in mice, but also across other animal species, including humans.

## Acknowledgements

We thank Amandine Cavaroc and Lisa Vierbaum for technical support; Henning Fenselau and Katrin Schrenk-Siemens for inspiring discussions and critical reading of the manuscript; Anke Tappe-Theodor for her help with corticosterone measurements; the Nikon Imaging Center at Heidelberg University for support with confocal microscopy. Funding: The authors gratefully acknowledge the data storage service SDS@hd supported by the Ministry of Science, Research and the Arts Baden-Württemberg (MWK) and the German Research Foundation (DFG) through grant INST 35/1314-1 FUGG. This work was supported by the European Research Council ERC-CoG-772395, the German research Foundation SFB/TRR 152 (to J.S.) and the European Molecular Biology Organization (EMBO) postdoctoral fellowship (to S.N.).

## Author Contributions

J.S. together with S.N. conceived the project; S.N. performed all experiments.

## Methods

### Mice

The following mouse lines were used in this study: LepR-cre (B6.129-Leprtm3(cre) Mgmj/J; The 920 Jackson Laboratory, IMSR Cat# JAX:032457), Vgat-FlpO (B6.Cg-Slc32a1tm1.1(flpo)Hze/J Cat# JAX:029591). By crossing LepR-Cre mice with Vgat-FlpO mice, we generated dual Cre/FlpO transgenic mice that, in combination with a Cre– and FlpO-dependent viral vector, allowed the selective targeting of POA vGat-negative or VGat-positive LepR neurons. Mice of either sex were used in experimental procedures. Mice were housed at room temperature (RT, 23 ± 1°C) in air-conditioned lab space / animal vivarium with a standard 12-h light/dark cycle and *ad libitum* access to food and water. All genetically modified mice in this study were on the C57BL/6N background.

All animal procedures were in accordance with the local ethics committee and governing body (Regierungspräsidium Karlsruhe, Germany) and were approved under protocol numbers: G-168/15, G-169/18; and G-181/2.

### Virus constructs. The following AAVs and titers were used

– ssAAV-DJ/2-hSyn1-chI-dlox-hChR2(H134R)_mCherry(rev)-dlox-WPRE-hGHp(A)AAV (titer: 5.3 x 10E12 vg/m, Zurich Viral Core)
– ssAAV-DJ/2-hSyn1-chI-dlox-mCherry(rev)-dlox-WPRE-hGHp(A) (titer: 7.2 x 10E12 vg/ml, Zurich Viral Core)
– ssAAV-retro/2-shortCAG-dlox-EGFP(rev)-dlox-WPRE-SV40p(A) (titer: 4.3 x 10E12 vg/ml, Zurich Viral Core)
– ssAAV-retro/2-CAG-dlox-tdTomato(rev)-dlox-WPRE-bGHp(A) (titer: 3.4 x 10E12 vg/ml, Zurich Viral Core)
– ssAAV DJ/2-hSyn1-chI-Con/Fon(hChR2(H134R)_EYFP)-WPRE-hGHp(A) (Cre-ON; FlpO-ON expression of ChR2; Zurich Viral Core, titer unknown)
– ssAAV DJ/2-hSyn1-CreON/FlpoOFF_ChR2_EYFP (Cre-ON; FlpO-OFF expression of ChR2; Zurich Viral Core, titer unknown)

### Stereotaxic virus injection and optical cannula implantation

Before surgery, 6-to 8-week-old mice were anesthetized using an intraperitoneal (i.p) injection of anesthesia mix (Medetomidine 0.5 mg/kg, Midazolam 5 mg/kg and Fentanyl 0.05 mg/kg). Mice were checked for the absence of the tail-pinch reflex as a sign of sufficient anesthesia. The mice were then immobilized in a stereotaxic frame (Model 1900; Kopf, USA) with ear bars (David Kopf Instruments), and ophthalmic ointment (Bepanthen; Bayer, Germany) was applied to prevent eye drying. The body temperature of animals was kept at 37°C using a heating pad.

After making an incision to the midline of the scalp, small unilateral craniotomies of approximately 0.6 mm diameter were performed with a hand drill (OS40; Osada Electric, Japan). The tips of glass capillaries (20–40 µm tip diameter) loaded with specific recombinant adeno-associated virus (ssAAV) carrying the functional construct or only the fluorescent protein were placed into specific brain areas.

The coordinates for target injection areas included the VMPO (AP, +0.75 mm; ML, −0.2 mm; DV, −4.8 mm from dura), the PVH (AP, −0.6 mm; ML, −0.26 mm; DV, −4.7 mm from dura), the DMH (AP, −1.4 mm; ML, −0.3 mm; DV, −4.9 mm from dura), and the VLPAG (AP, −4.72 mm; ML, −0.5 mm; DV, −2.10 mm from dura), as determined by the mouse brain atlas (Paxinos and Franklin Mouse Brain Atlas, 4th edition). A total of 250nl of virus-containing solution was injected unilaterally using a manual air pressure system. After injection, the capillary was left in place for an additional 5 min to allow for diffusion of the virus solution and then withdrawn.

Within the same AAV injection session, a 200μm diameter fiber optic probe (numeric aperture 0.53, Cat# FT200UMT, 1170 ThorLabs) was inserted to target the somas of preoptic LepR cells (in VMPO) or terminals (either in the PVH, the DMH, or the vlPAG). The coordinates used for the probe were identical to those used for the AAV injection, except for a 0.4mm upward adjustment in the dorsal-ventral position to ensure placement of the optic fiber above the area of infection. The probe was anchored to the skull with dental acrylic.

The scalp incision was closed with sterile absorbable-needled sutures (Marlin 17241041; Catgut, Germany), and mice received subcutaneous injection of Carprofen at 5mg/kg (Rimadyl; Zoetis, USA) for pain relief. Subsequently, anesthesia was antagonized with subcutaneous injection of Atipamezole at 2.5mg/kg, Flumazenil at 0.5mg/kg, and Naloxon at 1.2mg/kg. Mice were transferred to their home cages. For postoperative care, a second dose of Caprofen was administered after 24 hours. The mice cages were kept on a veterinary heating pad at 37°C for 12 hours and closely monitored. A minimum of 4 weeks was allowed for viral expression before any experiments were conducted.

### Optogenetic stimulation of LepR cells

Optogenetic stimulation experiments were performed in adult LepRCre mice, at least 4 weeks after AAV injection/implantation procedure.

To activate ChR2-expressing LepR neurons, a fiber optic probe was connected via an FC/PC adaptor to a 473-nm blue LED (Optogenetics-LED-Blue, Prizmatix). All experiments were conducted unilaterally, and the fiber optic cable was connected at least 2hr before the experiments to allow for habituation. During optogenetic probing, mice received light pulses with a power of 4–6 mW, a width of 10ms, and varying stimulation frequencies (5, 10, 20Hz) using a Prizmatix Pulser software and pulse train generator (Prizmatix). Each optogenetic probing session consisted of 1 minute of light stimulation followed by a 3 minute inter-stimulation interval.

Following the experiments, each mouse was euthanized (refer to procedure below), and the location of optical implant and viral expression and spread were examined under a fluorescence microscope. Mice with off-target implants, poorly expressed virus, or viral spread to other areas were excluded.

### Tail, BAT, core body and locomotor activity temperature recordings

Except for animals designated for electrophysiological recordings, all subjects received an i.p. injection of the anesthesia mix (Medetomidine 0.5mg/kg, Midazolam 5mg/kg and Fentanyl 0.05mg/kg). Following this, the abdominal fur was removed, skin disinfected with Braunol (Cat# 3864065, Braun, Germany), and cornea protected with Bepanthen ointment (Bayer, Germany). An implantable physiological signal wireless telemetry transmitter (TA11TA-F10; Data Sciences International, USA) was implanted in the abdominal cavity of a mouse, after which the muscle and skin layers were sutured separately with absorbable surgical threads (Marlin Cat#17241041, Catgut, Germany). After the surgery, the anesthesia was antagonized with Atipamezole at 2.5mg/kg, Flumazenil at 0.5mg/kg, and Naloxon at 1.2 mg/kg. Animals were monitored for recovery as outlined previously; a minimum recovery period of one week was observed before further procedures were undertaken.

The system consists of an implant, a receiver, a data converter (DEM), and a data analysis computer (IRBIS 3 software). The core body temperature signal was converted into a radio signal and received by a receiver (RSC-1, DSI, USA) positioned under the recording cage with a sampling rate of 5 min. Telemetry data were recorded using Ponemah 1162 software (DSI, USA).

For measuring tail temperatures and brown adipose tissue temperatures, we employed an infrared thermal camera (Vario CAM, InfraTec, Germany). Snapshot images were taken every 5 min using IRBIS 3 software (InfraTec, Germany). The average temperature was calculated at the midpoint of the tail (segment length of 1 cm) and at the center of the interscapular region, which was shaved 3 days prior to measurements.

### Food intake test

Food intake was assessed over a 12-hour period during the dark phase by quantifying pellet consumption. At the commencement of optogenetic stimulation, each mouse was housed in clear-bottom cages containing pre-weighed mouse chow pellets and a 1-inch layer of fresh sawdust bedding. Optogenetic stimulation was administered in cycles of 1 minute ON followed by 3 minutes OFF, with a stimulation frequency of 10Hz, spanning a 12-hour duration (from 7AM to 7PM). At the end of the stimulation, both the consumed pellet weight and the mouse’s body weight were measured.

### Fecal corticosterone metabolite measurements

Corticosterone metabolite measurements were carried out on fecal samples as previously described (Touma et al. Gen Comp Endocrinol, 2003; DOI: 10.1016/s0016-6480(02)00620-2.) with slight modifications as described in Segeleck et al. (Sci Rep. 2023; DOI: 10.1038/s41598-023-29052-7.). In brief, feces were collected from controls (mCherry injected) and ChR2-positive animals around 7AM, directly after the overnight optogenetic stimulation cycle. Fecal boli (5–6 fecal boli per animal) were stored at −20°C. Fecal samples were dried for two hours at 80°C before mechanical homogenization, and 0.05g were extracted with 1ml 80% methanol for 30min on a vortex. After centrifugation for 10min at 2500g, 0.5ml supernatant was frozen until analysis. Fecal corticosterone metabolite was measured using a 5α-pregnane-3β,11β,21-triol-20-one enzyme immunoassay (EIA).

### Electrophysiology and photo-stimulation in brain slices

Four weeks after surgery, the mice were deeply anesthetized with Ketamine/Xylaxine mixture (Ketamine: 220mg/kg, Ketavet; Zoetis, USA and Xylazine 16mg/kg, Rompun; Bayer, Germany) and decapitated. Brains were dissected quickly and chilled in ice-cold (4°C) artificial CSF (aCSF) containing the following (in mM): NaCl, 85; KCl, 2.5; glucose, 10; sucrose, 75; NaH2PO4, 1.25; NaHCO3, 25; MgCl, 3; CaCl2, 0.1; myo-inositol, 3; sodium pyruvate, 2; ascorbic acid, 0.4. Coronal slices (250μm thick) of MnPOA were prepared using a vibratome (Leica VT1200S, Germany) and then incubated for 7 minutes in heated (32°C) oxygenated holding aCSF (in mM): NaCl, 109; KCl, 4; glucose, 35; NaH2PO4, 1.25; NaHCO3, 25; MgCl2, 1.3; CaCl2, 1.5. Individual slices were then transferred to the recording chamber where they were continuously superfused with oxygenated recording aCSF at ∼2 ml/min.

Neuronal action potentials in LepR cells were recorded at room temperature in whole-cell current-clamp configuration, with borosilicate glass patch pipette (4–8MΩ resistance) filled with internal solution containing the following (in mM): K-gluconate, 138; KCl, 1369 2; NaCl, 5; HEPES, 10; EGTA, 10 (or equimolar amount of BAPTA); CaCl2, 1; Mg-ATP, 1. Micropipettes (O.D. 1.5 mm, I.D. 0.86 mm, Sutter Instrument, BF150-86-7.5) were pulled on a micropipette puller (P-97, Sutter Instrument, USA).

Intracellular solution was passed through 0.22µm filter before filling the electrode pipette. The open pipette resistance was between 4-8 MΩ.

Cells in acute POA slices were visualized using a SliceScope upright microscope (Scientifica, UK) equipped with a 40X water immersion objective (U-TV1X-2, Olympus, Japan). Images were acquired by a digital CCD camera (ORCA-R2 C10600-10B, Hamamatsu Photonics K.K., Japan) using Micro Manager 1.4 software (Vale’s lab, UCSF, USA). Electrophysiological recordings were acquired using a MultiClamp 700B amplifier (Molecular Devices, USA), together with an Axon Digidata 1550B digitizer (Molecular Devices, USA) and Clampex 11.0.3 software (Molecular Devices, USA). All signals were sampled at 20kHz and low pass filtered at 10kHz.

To illuminate neurons infected with channel-rhodopsin within the POA, blue light (470 nm) pulses were applied by a light-emitting diode (LED)-based optical system (pE-100 CoolLed, Scientifica, UK). The target site was illuminated with 10ms light pulses at variable stimulation frequency (5, 10, and 20Hz). Each train of the light pulses was given for 60s.

### Histology of AAV-injected mouse brains

Mice were anesthetized, transcardially perfused with paraformaldehyde (PFA), and then decapitated. The whole heads were immersed in 4% PFA for at least one day at 4°C. Subsequentially, the brains were removed from the skull and transferred to a phosphate-buffered saline (PBS) solution containing sucrose. Coronal sections of 30µm were cut at the microtome and stored at –20°C in cryoprotectant solution. These brain sections were later stained for GFP/mCherry according to the procedure outlined below.

### Immunohistochemistry procedure

Animals were deeply anesthetized with isoflurane and transcardially perfused with a phosphate buffer solution (PBS; 3.85 g of NaOH and 16.83 g of NaH2PO4 in 1l of distilled water) followed by a 4% paraformaldehyde (PFA) solution. Brains were dissected out and left overnight (O/N) in 4% PFA at 4°C. Over the following 2 days brains were immersed into PBS/sucrose solutions (24 hours in 10% sucrose followed by 30% sucrose, until brains sank to the bottom of container tube). Brains were sectioned with a cryo-microtome at 30μm thickness and sections (free-floating) were kept in cryo-protectant solution (250ml glycerol; 250ml ethylene glycol filled up with PBS to 1l) at 4°C until mounting. Tissue was washed extensively with PBSX 0.1 and once with PBS after which sections were mounted using Immu-Mount (Cat# 9990402, Fisher Scientific, UK) onto glass slides. Confocal images were taken at the Nikon imaging center at Heidelberg University, with the Nikon A1R confocal microscope under Nikon Plan Apo λ 10x magnification NA 0.45 (working distance 4mm, the field of view 1.27 x 1.27 mm) objective.

### Retrogradely labeled LepR neurons

For the retrograde tracing of POA LepR neurons, we used the LepR-*Cre* mice and injected the PVH and DMH with retro-AAV (Viral Vector Facility from the University of Zurich; Switzerland) conjugated with Cre-dependent EGFP or tdTomato (see above for specification), respectively (250nl, unilateral). Confocal microscopic images of the POA area were taken at the Nikon imaging center at Heidelberg University with the Nikon 989 A1R confocal microscope under Nikon Plan Apo λ 10x magnification NA 0.45 (working distance 990 4mm, the field of view 1.27 x 1.27 mm) objective. For each mouse, images of 3-4 representative fields of the POA hypothalamic area were acquired. Double-projecting POA neurons were identified based on cell bodies labeled with both tdTomato and EGFP. The proportion of double projecting POA neurons to POA neurons projecting to either the PVH or DMH was also calculated for each mouse. Images presented were processed with ImageJ.

### Statistical Analysis

Statistical analysis was performed using GraphPad Prism 6 (GraphPad Software, San Diego, CA). Data were expressed as mean ± SEM and were analyzed using one-way or two-way ANOVA, as indicated. Dunnett’s multiple comparisons test was used as post hoc comparisons of ANOVA. A value of p<0.05 was considered statistically significant.

## Supplementary Figures

**Supplementary Figure 1.**
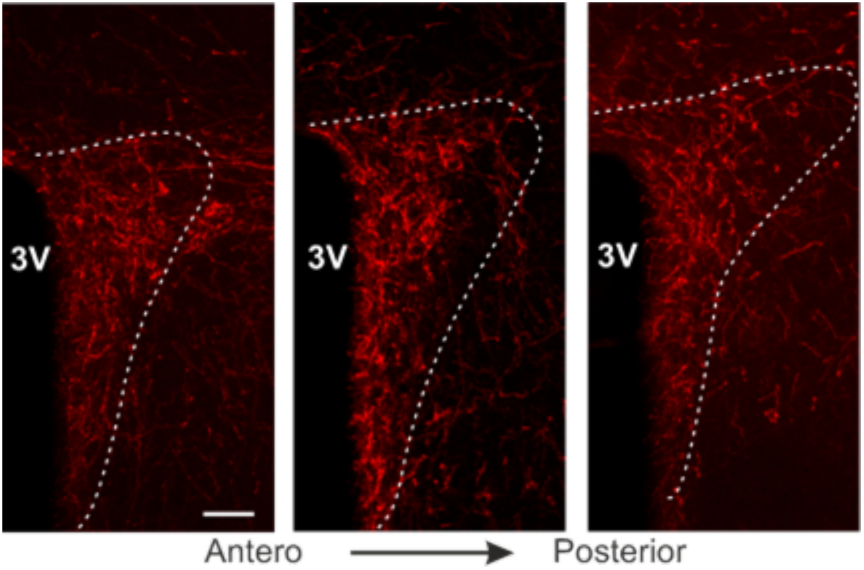
– LepR neurons densely innervate the PVH. Representative histological images showing the projections of VMPOLepR neurons (indicated in red, ChR2-mCherry) in the PVH area (anterior to posterior). 3V, third ventricle. Scale bar: 50um.

**Supplementary Figure 2.**
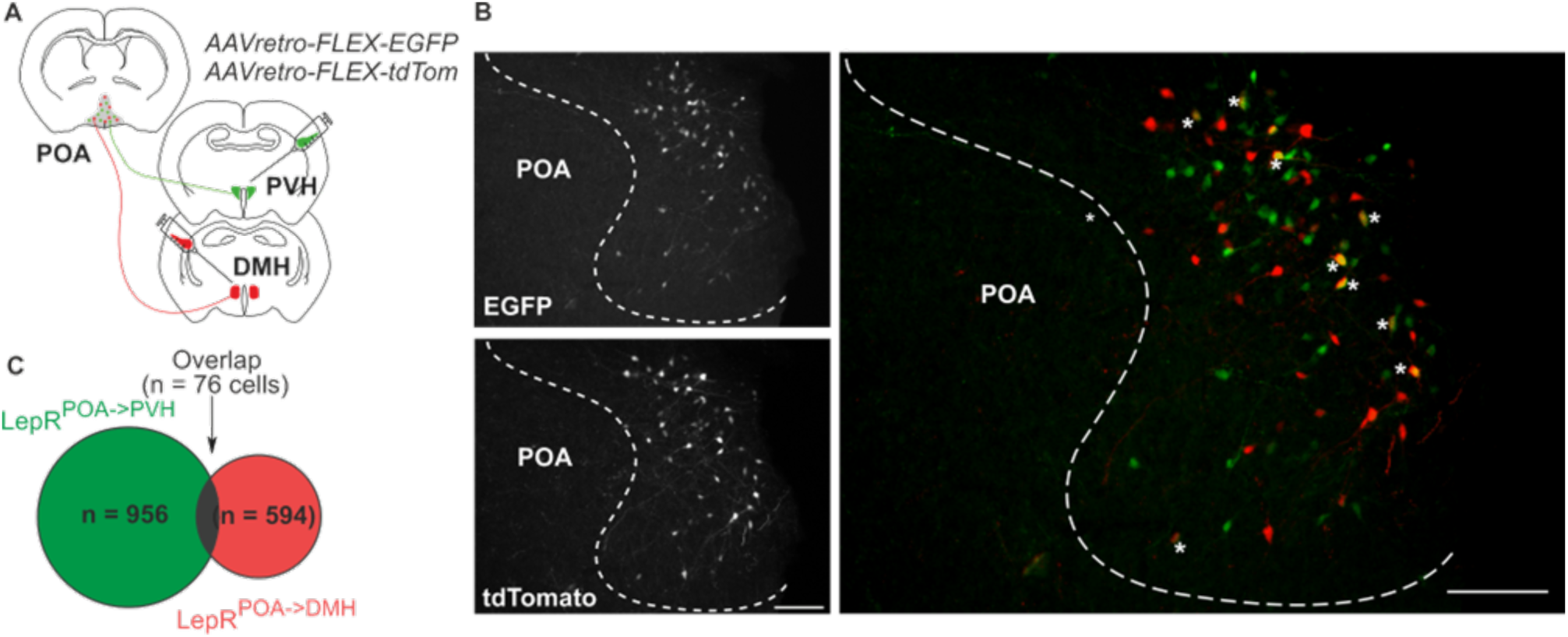
– VMPO^LepR^ neurons projecting to PVH and DMH are largely non-overlapping. **A**. Strategy used for labeling of VMPO^LepR^ neurons that project to either PVH or DMH. ssAAVretro-EGFP and ssAAVretro-tdTomato were injected in PVH and DMH regions, respectively. **B.** Fluorescence distribution of EGFP (from AAV-retro particles injected into the PVH) and tdTomato (from AAV-retro particles injected into the DMH) expression in POA LepR projection neurons. Scale bar: 100µm. **C.** Pie chart depicting the number of VMPO^LepR^ neurons in the preoptic area (POA) that are positive for EGFP (projecting to PVH), tdTomato (projecting to DMH), and double-labeled neurons that project to both areas.

**Supplementary Figure 3.**
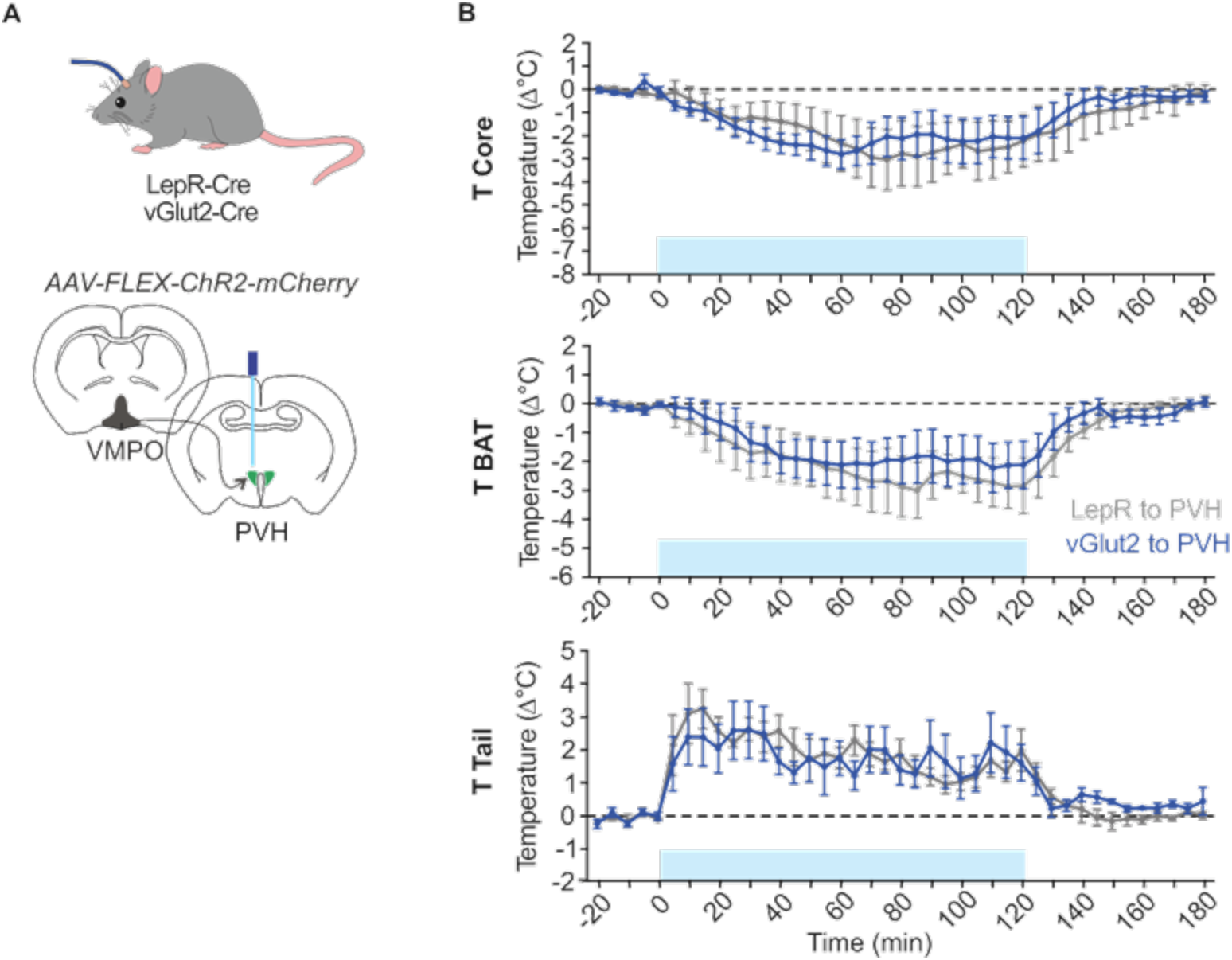
– VMPO^VGlut2^ neuronal projections to the PVH mirror T_core_, T_BAT_ and T_tail_ responses observed for VMPO^LepR^ neuronal projections to the PVH. **A.** Schematic representation of the optogenetic experiment: Cre-dependent ChR2 AAV virus was injected into the VMPO of either vGlut2-Cre or LepR-Cre mice, with the optogenetic probe subsequently implanted in the downstream PVH projection area. **B.** Optogenetic stimulation at 10Hz of VMPO^VGlut^^2^ neuron terminals in the PVH of ChR2-positive VGlut2-Cre animals (N = 5 mice) induced a decrease in body core (upper panel) and BAT (middle panel) temperature, as well as an increase in tail temperature (bottom panel), similar to results obtained for PVH-projecting of VMPO^LepR^ neuron terminals (N = 5 mice, same data as shown in Figure 3D).

## References

1 Gordon, C. J. Temperature Regulation in Laboratory Rodents., (Cambridge University Press, 1993).

2 International Symposium on the Nutrition of, H., Hacker, J. B., Ternouth, J. H., Australian Society of Animal, P. & International Symposium on the Nutrition of, H. The Nutrition of herbivores. (Academic Press, 1987).

3 Marriott, B. M. & Institute of Medicine (U.S.). Committee on Military Nutrition Research. Nutritional needs in hot environments: applications for military personnel in field operations. (National Academy Press, 1993).

4 Horvath, T. L., Stachenfeld, N. S. & Diano, S. A temperature hypothesis of hypothalamus-driven obesity. Yale J Biol Med 87, 149–158 (2014).

5 Morrison, S. F. & Nakamura, K. Central Mechanisms for Thermoregulation. Annu Rev Physiol 81, 285–308 (2019). 10.1146/annurev-physiol-020518-114546

6 Morrison, S. F., Nakamura, K. & Madden, C. J. Central control of thermogenesis in mammals. Exp Physiol 93, 773–797 (2008). 10.1113/expphysiol.2007.041848

7 Tran, L. T. et al. Hypothalamic control of energy expenditure and thermogenesis. Exp Mol Med 54, 358–369 (2022). 10.1038/s12276-022-00741-z

8 An, J. J., Liao, G. Y., Kinney, C. E., Sahibzada, N. & Xu, B. Discrete BDNF Neurons in the Paraventricular Hypothalamus Control Feeding and Energy Expenditure. Cell Metab 22, 175–188 (2015). 10.1016/j.cmet.2015.05.008

9 Houtz, J., Liao, G. Y., An, J. J. & Xu, B. Discrete TrkB-expressing neurons of the dorsomedial hypothalamus regulate feeding and thermogenesis. Proc Natl Acad Sci U S A 118 (2021). 10.1073/pnas.2017218118

10 Schneeberger, M. et al. Regulation of Energy Expenditure by Brainstem GABA Neurons. Cell 178, 672–685 e612 (2019). 10.1016/j.cell.2019.05.048

11 Yu, S. et al. Glutamatergic Preoptic Area Neurons That Express Leptin Receptors Drive Temperature-Dependent Body Weight Homeostasis. J Neurosci 36, 5034–5046 (2016). 10.1523/JNEUROSCI.0213-16.2016

12 Siemens, J. & Kamm, G. B. Cellular populations and thermosensing mechanisms of the hypothalamic thermoregulatory center. Pflugers Arch 470, 809–822 (2018). 10.1007/s00424-017-2101-0

13 Tan, C. L. & Knight, Z. A. Regulation of Body Temperature by the Nervous System. Neuron 98, 31–48 (2018). 10.1016/j.neuron.2018.02.022

14 Madden, C. J. & Morrison, S. F. Central nervous system circuits that control body temperature. Neurosci Lett 696, 225–232 (2019). 10.1016/j.neulet.2018.11.027

15 Jais, A. & Bruning, J. C. Arcuate Nucleus-Dependent Regulation of Metabolism-Pathways to Obesity and Diabetes Mellitus. Endocr Rev 43, 314–328 (2022). 10.1210/endrev/bnab025

16 Bruning, J. C. & Fenselau, H. Integrative neurocircuits that control metabolism and food intake. Science 381, eabl7398 (2023). 10.1126/science.abl7398

17 Atasoy, D., Betley, J. N., Su, H. H. & Sternson, S. M. Deconstruction of a neural circuit for hunger. Nature 488, 172–177 (2012). 10.1038/nature11270

18 Fenselau, H. et al. A rapidly acting glutamatergic ARC-->PVH satiety circuit postsynaptically regulated by alpha-MSH. Nat Neurosci 20, 42–51 (2017). 10.1038/nn.4442

19 Garfield, A. S. et al. A neural basis for melanocortin-4 receptor-regulated appetite. Nat Neurosci 18, 863–871 (2015). 10.1038/nn.4011

20 Li, M. M. et al. The Paraventricular Hypothalamus Regulates Satiety and Prevents Obesity via Two Genetically Distinct Circuits. Neuron 102, 653–667 e656 (2019). 10.1016/j.neuron.2019.02.028

21 Madden, C. J. & Morrison, S. F. Neurons in the paraventricular nucleus of the hypothalamus inhibit sympathetic outflow to brown adipose tissue. Am J Physiol Regul Integr Comp Physiol 296, R831–843 (2009). 10.1152/ajpregu.91007.2008

22 Kong, D. et al. GABAergic RIP-Cre neurons in the arcuate nucleus selectively regulate energy expenditure. Cell 151, 645–657 (2012). 10.1016/j.cell.2012.09.020

23 Qian, S. et al. A temperature-regulated circuit for feeding behavior. Nat Commun 13, 4229 (2022). 10.1038/s41467-022-31917-w

24 Suwannapaporn, P., Chaiyabutr, N., Wanasuntronwong, A. & Thammacharoen, S. Arcuate proopiomelanocortin is part of a novel neural connection for short-term low-degree of high ambient temperature effects on food intake. Physiol Behav 245, 113687 (2022). 10.1016/j.physbeh.2021.113687

25 Friedman, J. M. Leptin and the endocrine control of energy balance. Nat Metab 1, 754–764 (2019). 10.1038/s42255-019-0095-y

26 Fischer, A. W. et al. Leptin Raises Defended Body Temperature without Activating Thermogenesis. Cell Rep 14, 1621–1631 (2016). 10.1016/j.celrep.2016.01.041

27 Machado, N. L. S. & Saper, C. B. Genetic identification of preoptic neurons that regulate body temperature in mice. Temperature, 1–9 (2021). 10.1080/23328940.2021.1993734

28 Abbott, S. B. G. & Saper, C. B. Median preoptic glutamatergic neurons promote thermoregulatory heat loss and water consumption in mice. J Physiol 595, 6569–6583 (2017). 10.1113/JP274667

29 Kroeger, D. et al. Galanin neurons in the ventrolateral preoptic area promote sleep and heat loss in mice. Nat Commun 9, 4129 (2018). 10.1038/s41467-018-06590-7

30 Song, K. et al. The TRPM2 channel is a hypothalamic heat sensor that limits fever and can drive hypothermia. Science 353, 1393–1398 (2016). 10.1126/science.aaf7537

31 Tan, C. L. et al. Warm-Sensitive Neurons that Control Body Temperature. Cell 167, 47–59 e15 (2016). 10.1016/j.cell.2016.08.028

32 Andermann, M. L. & Lowell, B. B. Toward a Wiring Diagram Understanding of Appetite Control. Neuron 95, 757–778 (2017). 10.1016/j.neuron.2017.06.014

33 Hao, S. et al. The Lateral Hypothalamic and BNST GABAergic Projections to the Anterior Ventrolateral Periaqueductal Gray Regulate Feeding. Cell Rep 28, 616–624 e615 (2019). 10.1016/j.celrep.2019.06.051

34 Imoto, D. et al. Refeeding activates neurons in the dorsomedial hypothalamus to inhibit food intake and promote positive valence. Mol Metab 54, 101366 (2021). 10.1016/j.molmet.2021.101366

35 Morrison, S. F. & Nakamura, K. Central neural pathways for thermoregulation. Front Biosci (Landmark Ed) 16, 74–104 (2011). 10.2741/3677

36 Zhang, K. X. et al. Violet-light suppression of thermogenesis by opsin 5 hypothalamic neurons. Nature 585, 420–425 (2020). 10.1038/s41586-020-2683-0

37 Hrvatin, S. et al. Neurons that regulate mouse torpor. Nature 583, 115–121 (2020). 10.1038/s41586-020-2387-5

38 Takahashi, T. M. et al. A discrete neuronal circuit induces a hibernation-like state in rodents. Nature 583, 109–114 (2020). 10.1038/s41586-020-2163-6

39 Zhang, Z. et al. Estrogen-sensitive medial preoptic area neurons coordinate torpor in mice. Nat Commun 11, 6378 (2020). 10.1038/s41467-020-20050-1

40 Swoap, S. J. The pharmacology and molecular mechanisms underlying temperature regulation and torpor. Biochem Pharmacol 76, 817–824 (2008). S0006-2952(08)00384-5 [pii] 10.1016/j.bcp.2008.06.017

41 Upton, B. A., D’Souza, S. P. & Lang, R. A. QPLOT Neurons-Converging on a Thermoregulatory Preoptic Neuronal Population. Front Neurosci 15, 665762 (2021). 10.3389/fnins.2021.665762

42 Krashes, M. J. et al. An excitatory paraventricular nucleus to AgRP neuron circuit that drives hunger. Nature 507, 238–242 (2014). 10.1038/nature12956

43 Sutton, A. K. et al. Control of food intake and energy expenditure by Nos1 neurons of the paraventricular hypothalamus. J Neurosci 34, 15306–15318 (2014). 10.1523/JNEUROSCI.0226-14.2014

44 Gavrilova, O. et al. Torpor in mice is induced by both leptin-dependent and – independent mechanisms. Proc Natl Acad Sci U S A 96, 14623–14628 (1999). 10.1073/pnas.96.25.14623

45 Yu, S. et al. Preoptic leptin signaling modulates energy balance independent of body temperature regulation. Elife 7 (2018). 10.7554/eLife.33505

